# Harnessing Coding Sequence Cleavage: Theophylline Aptazymes as Portable Gene Regulators in Bacteria

**DOI:** 10.1101/2025.04.24.650125

**Authors:** Emre Yurdusev, Yasmine Nicole, Felicia Pan Du, Nahide Ekin Suslu, Jonathan Perreault

## Abstract

Nucleic acid-based regulatory elements capable of modulating gene expression in response to specific molecular cues have gained increasing attention in synthetic biology. These systems, which include riboswitches, allosteric DNAzymes, and aptazymes, function as Gene Expression Nucleic Allosteric actuators (GENAs) by coupling molecular recognition with genetic regulation. Their versatility can enable applications in diagnostics, therapeutics, and metabolic engineering. This study presents a novel “semi-trans” aptazyme-based system for gene regulation in bacteria, expanding the range of GENAs. The system employs theophylline-responsive hammerhead ribozyme aptazymes positioned in the 5’ UnTranslated Region (UTR), designed to cleave within the coding sequence of the target gene, thereby modulating gene expression in a ligand-dependent manner. Using the *tetA* gene in *Escherichia coli* (*E. coli*) as a proof of concept, we demonstrate ligand-controlled regulation of tetracycline resistance and nickel sensitivity. The system′s effectiveness is validated through in vitro cleavage assays and in vivo phenotypic studies in two *E. col*i strains, highlighting its portability across genetic backgrounds. Furthermore, the ability to design multiple aptazymes targeting different coding regions enables complex and fine-tuned regulation. This work broadens the landscape of synthetic gene regulation tools, facilitating the development of new aptazymes based on this approach.

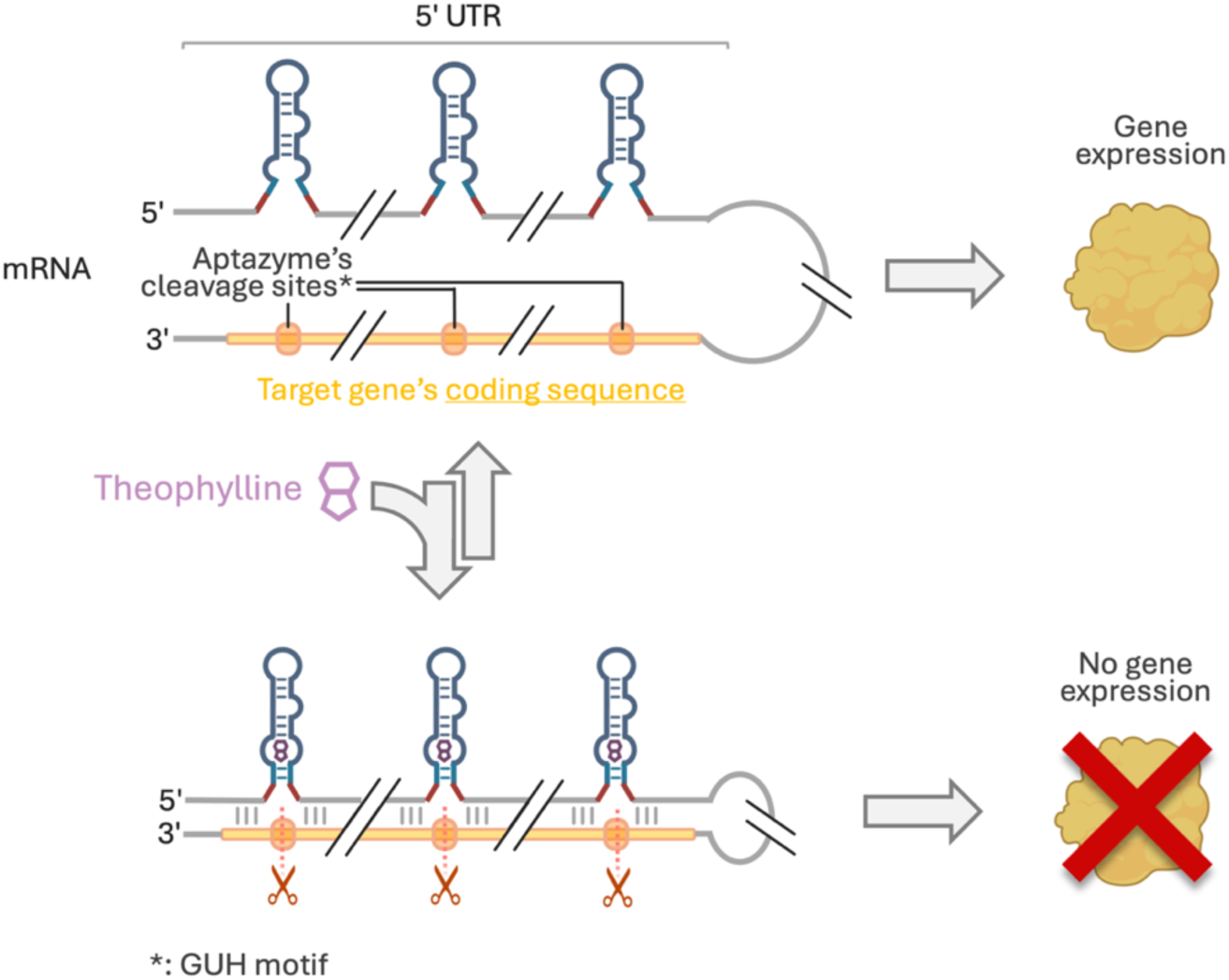

## Introduction

Aptazymes, RNA fragments with catalytic activity modulated by specific ligand binding, have emerged as versatile tools for synthetic biology ^1^. As allosteric regulators of gene expression, aptazymes offer an unprecedented ability to link the presence of environmental molecules to the precise modulation of genetic activity. Similar tools, such as [2] riboswitches ^2,3^, which regulate gene expression without necessarily relying on enzymatic activity, collectively highlight the potential of Gene Expression Nucleic Allosteric actuators (GENAs) in diagnostics, therapeutics, and metabolic engineering. These systems capitalize on their ability to couple molecular recognition with gene expression regulation. Striking examples include riboswitches engineered for microbial metabolic engineering, which dynamically regulate biosynthetic pathways to optimize metabolite production ^4,5^. For instance, engineered riboswitches in *Corynebacterium glutamicum* have been used to modulate lysine biosynthesis, showcasing their potential for industrial biotechnology applications ^4^. Similarly, allosteric DNAzymes ^6^—the DNA counterparts of RNA aptazymes—have been integrated into biosensors for highly sensitive biomarker detection, contributing to early cancer diagnostics ^7^. The application of ligand-responsive GENAs in viral vectors has further demonstrated their potential to provide dynamic and reversible control over therapeutic gene expression, reinforcing their relevance in gene therapy and synthetic biology. Aptazymes, for example, have also been successfully applied for gene silencing and therapeutic control, including their incorporation into RNA-virus-based episomal vectors (REVec) for regulated gene expression ^8^. In this system, aptazymes inserted into the UTRs of transgenes (e.g., GFP, NanoLuc) or essential viral genes (e.g., phosphoprotein P) enable ligand-dependent RNA cleavage, leading to controlled gene suppression. Beyond gene regulation, this strategy allows vector elimination when aptazymes target genes critical for REVec replication. Specifically, ligand-induced cleavage of the P gene mRNA prevents production of a key replication protein, progressively reducing vector levels over time. Since REVec cannot reinfect cells, prolonged ligand exposure results in complete clearance of the vector from transduced cells. This provides a built-in biocontainment mechanism, enhancing safety in gene therapy applications. Expanding on this principle, GENAs have also been integrated into viral gene therapy platforms, such as adeno-associated viruses (AAVs) and oncolytic adenoviruses, to enable external control of transgene expression ^9–11^. These systems offer a precise alternative to traditional inducible promoters, increasing the number of trigger molecules that can be used to improve safety and tunability in therapeutic gene delivery ^11^.

Furthermore, GENAs have been incorporated into CRISPR technologies, significantly enhancing precision in therapeutic applications ^12^. By embedding ligand-responsive GENAs into guide RNAs (gRNAs), researchers have developed CRISPR systems that activate or suppress Cas9 activity in response to specific small molecules or environmental cues ^13,14^. This approach enables fine-tuned gene editing control, increasing both the safety and specificity of applications such as targeted gene therapies ^12^. Beyond gene editing, GENA-CRISPR platforms have been incorporated into biosensing and diagnostics, facilitating the development of highly sensitive and selective systems for biomarkers, pathogens, and environmental detection, further broadening their utility beyond therapeutics ^15^. Through these applications, GENAs establish themselves as a powerful and modular framework for engineering gene regulation, with implications across diverse fields from metabolic engineering to precision medicine.

The ongoing goals in GENA-based gene regulation are twofold: expanding the portfolio of GENAs to respond to a broader range of molecules and refining their mechanisms for more efficient and translatable gene regulation. To address the first goal, methods such as appending existing aptamers to ribozyme cores or expression platforms have been developed. These designs use either rational approaches ^16^ or random linker sequences ^17,18^ followed by SELEX (Systematic Evolution of Ligands by Exponential Enrichment) to identify functional GENAs. Additionally, SELEX has been employed to generate entirely synthetic GENAs, evolving both the aptameric and catalytic components for specific functionalities ^19–24^. This approach is particularly effective for aptazymes, as their cleavage activity provides a direct, selectable readout compatible with both in-vitro and in-vivo selection, facilitating high-throughput screening ^19–21^.

While increasing the diversity of GENAs is crucial, improving their efficiency and regulatory performance is equally important. This challenge is closely tied to the mechanisms by which GENAs regulate gene expression ^25,26^. Different classes of GENAs employ distinct modes of action: riboswitches modulate gene expression by influencing transcription termination ^27^, translation initiation ^3^, or RNA stability ^28^, depending on their interaction with target molecules. In contrast, aptazymes ^1,29^ and DNAzymes ^30^ primarily act through catalytic cleavage, disrupting critical mRNA structures to control gene expression. Among aptazymes, those based on hammerhead ribozymes (HHRs) are particularly well-suited for engineered gene regulation due to their compact catalytic core ^31^ and modular structure ^32,33^, allowing for precise control over cleavage sites, particularly at NUH motifs, to regulate gene expression efficiently ^34^.

Optimizing the placement of aptazyme elements or its site of action within target mRNAs further enhances regulatory efficiency. In practice, most aptazyme-based systems place the catalytic element in untranslated regions (UTRs) of the target mRNA. For example, aptazymes in the 3′ UTR may disrupt a stabilizing structure ^35,36^. Similarly, aptazymes in the 5′ UTR can regulate gene expression by cleaving to release a sequestered ribosome binding site (RBS) such as the Shine-Dalgarno sequence in bacterial system ^37^ or cell-free context ^38^ or altering mRNA stability ^36^. However, translation mechanisms and UTR functions differ between bacteria and eukaryotes, limiting portability. Moreover, cleavage effects can be context-dependent — sometimes increasing, other times decreasing gene expression — due to structural or stability changes unrelated to the cleavage event itself ^39–41^. Notably, several challenges in this field are documented in graduate theses rather than peer-reviewed articles, reflecting how often these efforts remain unpublished due to technical difficulties or limited success ^39,41^. Although some systems report promising results, the broader applicability of UTR-targeted aptazymes is still unpredictable and often challenging to reproduce. In this context, exploring aptazyme integration directly within coding sequences may offer a valuable alternative, potentially overcoming some of the limitations associated with UTR-based approaches. Recent work by Zhou *et al.*^34^ demonstrated the feasibility of HHR aptazymes targeting coding sequences. In bacterial models, the aptazyme was placed immediately downstream of the AUG start codon, while trans-acting designs effectively regulated gene expression in mammalian cells. This approach offers a high degree of portability, as it can be applied to virtually any gene in diverse organisms, regardless of promoter characteristics or UTR structures. The universality stems from the ubiquitous nature of coding sequences and the unequivocal consequence of mRNA cleavage within the coding sequence in all species, i.e. inability to translate the complete protein.

In our study, we propose a novel system using an aptazyme positioned in the 5′ UTR of the target mRNA, with theophylline-responsive HHR as a model. Acting in a semi-trans manner, i.e. within the same RNA molecule but from a long distance, this aptazyme is designed to cleave within the coding sequence, balancing the advantages of both cis and trans approaches. This strategy addresses challenges associated with cis-acting aptazymes, that are difficult to incorporate within coding regions without affecting the amino acid sequence, and avoids the complexities of long-range trans interactions with multiple promoters and expression systems. By positioning the aptazyme close to its target site, our design presumably enhances regulatory efficiency and minimizes unintended effects.

To validate this approach, we targeted the *tetA* gene in *E. coli* as a proof of concept. The *tetA* gene, known for its dual phenotype, conferring tetracycline resistance and nickel sensitivity ^42^. We demonstrated effective gene regulation through phenotypic changes, namely theophylline-mediated sensitivity to tetracycline and resistance to nickel. Additionally, we explored the additive effects of multiple aptazyme systems and assessed the correlation between in vitro and in vivo results.

## Results

### Aptazyme Design Strategy

To validate the effectiveness and universality of our aptazyme-based gene regulation approach, we selected the *tetA* gene as our target. The *tetA* gene provides a unique opportunity due to its dual phenotypical effects on bacteria ^42^: it confers resistance to tetracycline while simultaneously increasing sensitivity to nickel (**Figure 1A**). This enables easy selection of ligand-responsive GENAs through opposing survival pressures ^42^. For example, to isolate GENAs that downregulate *tetA* in response to a ligand, selection can alternate between tetracycline survival in the presence of the ligand and nickel survival in its absence. Conversely, selecting GENAs that downregulate *tetA* in the absence of the ligand and restore expression with the ligand involves reversing the survival conditions. As demonstrated by Murakana *et al* ^42^, this dual selection simplifies the identification of functional GENAs in *in vivo* context with precise ligand-dependent regulation. By establishing the proof of concept that our approach can phenotypically regulate *tetA*, we aim to achieve dual validation: demonstrating the efficacy of our system and showcasing its potential applicability in “*in vivo* SELEX” to identify new coding sequence-targeting semi-trans acting aptazymes.

**Figure 1:**
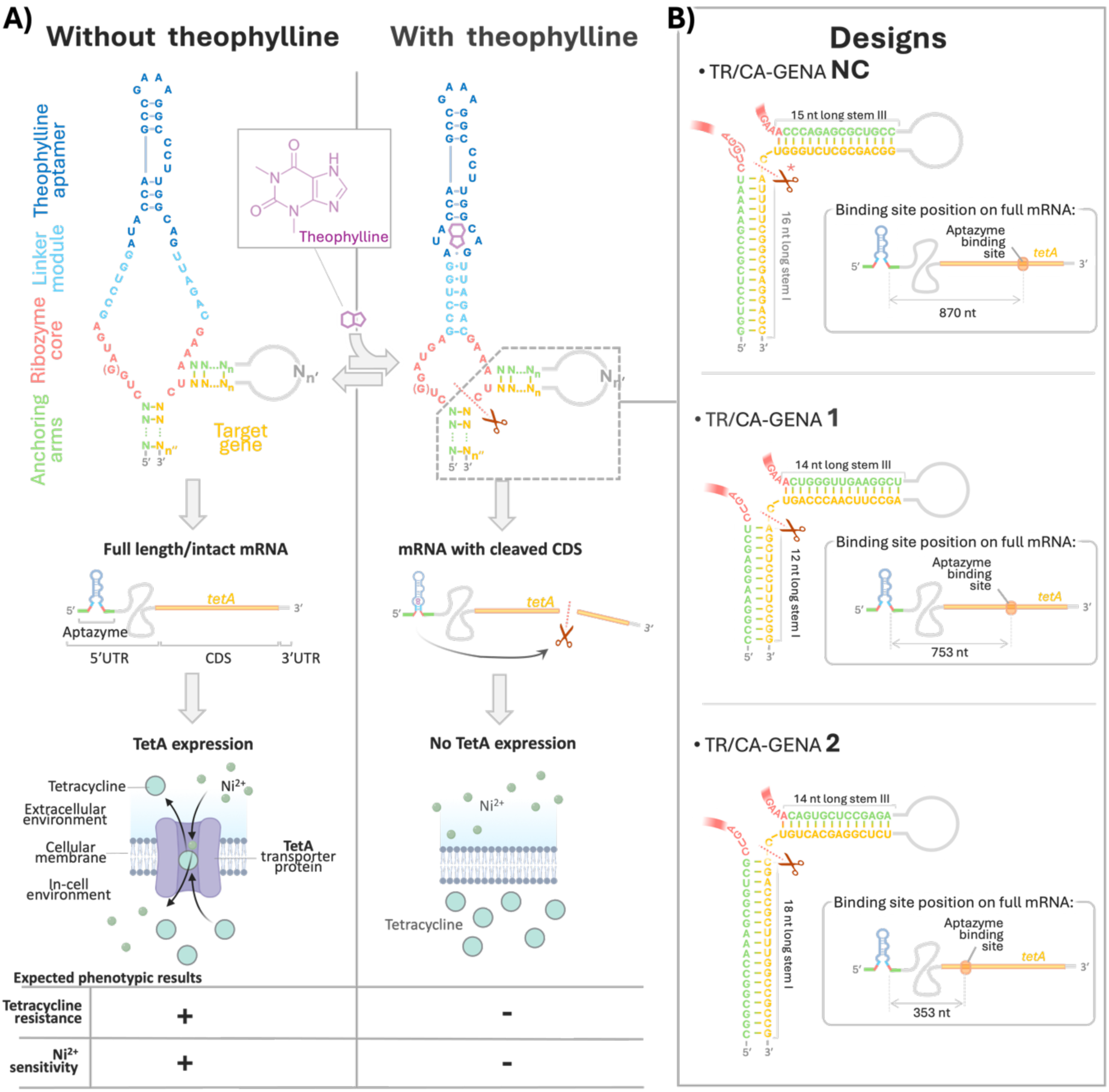
Theophylline-Responsive Aptazyme Regulation of TetA Expression in *E. coli.* **A)** The design and function of a theophylline-responsive hammerhead ribozyme (HHR) aptazyme system regulating *tetA* gene expression in *E. coli*. In the absence of theophylline, the aptazyme remains inactive, allowing full-length *tetA* mRNA translation and tetracycline resistance. Upon theophylline binding, the aptazyme is activated and cleaves the *tetA* mRNA coding sequence. Bottom: phenotypic effects of aptazyme activation, with expected bacterial growth outcomes under tetracycline selection in the presence or absence of theophylline. **B)** Aptazyme designs TR/CA-GENA NC, 1 and 2 are depicted with their different target sites within the tetA coding sequence. TR/CA-GENA NC contains an additional guanine (G) nucleotide within the ribozyme core (shown in brackets), which impairs its cleavage activity; accordingly, an asterisk besides the scissor symbol indicates impaired cleavage.

The aptazymes targeting *tetA* were designed using the theophylline-responsive hammerhead ribozyme (HHR) core, as described by Soukup *et al.* ^17^ and further characterized by Zhou *et al.* ^34^. Specifically, we employed a type III theophylline HHR aptazyme, characterized by an open stem III, which facilitates trans-acting configurations. The design strategy involved creating “half” aptazymes, where the functional ribozyme structure requires hybridization with the target gene to complete the catalytic core (**Figure 1A**). This approach ensures that the 5′ and 3′ arms of the aptazyme hybridize with sequences flanking a target site, with the required GUH motif (where H is A/C/U), within the coding sequence of *tetA* [43]. The missing segment of the ribozyme catalytic core is thus reconstituted only upon target binding, ensuring specificity.

For the design of these aptazymes, we utilized RiboSoft 2.0 software, developed by Kharma *et al.* ^43^, which implements a hammerhead ribozyme design algorithm. We selected three distinct designs with high scores, long hybridizing arms, and targeting different regions of the *tetA* coding sequence. Two distinct aptazyme designs (TR/CA-GENA 1, and 2) were developed, each targeting different sites within the *tetA* coding sequence. The acronym TR/CA-GENA stands for *Theophylline-Responsive, Coding-sequence-Acting Gene Expression Nucleic Acid*, reflecting both the ligand-dependent activation and the design’s intended location within coding regions. The binding site positions of each aptazyme within the *tetA* mRNA are depicted in Figure 1B, highlighting the modular nature of the system and its potential for multiplexed regulation. In designing these aptazymes, we maintained the integrity of the ribozyme catalytic core and the theophylline aptamer, as these elements are essential for ligand-dependent activation. Modifications were restricted to the stem regions, which are known to tolerate changes without disrupting the catalytic function.

In addition to the two functional cleaving designs, we also included a control aptazyme, TR/CA-GENA NC (NC for *Not Cleaving*), in which an extra guanine (G) was introduced into the ribozyme catalytic core (**Figure 1**). While this design retains the theophylline aptamer and can still bind to its specific target site, the ribozyme core is expected to be catalytically inactive or strongly impaired due to this disruption. This non-cleaving construct was incorporated to evaluate the importance of cleavage in the gene regulation process—specifically, to assess the extent to which target binding alone, in the absence of cleavage, contributes to regulatory outcomes.

We generated three distinct aptazyme designs, each targeting a different region within the *tetA* coding sequence. This aimed to demonstrate that: i) targeting coding sequences provides sufficient space for designing multiple aptazymes, ii) the ability to target different sites within the same gene illustrates the adaptability of the approach to diverse sequence fragments and gene types and iii) simultaneous use of multiple aptazymes can potentially amplify regulatory effects. To assay the latter, we thus also constructed combinations of the three aptazymes.

### In Vitro Testing

To evaluate the performance of the designed aptazymes, we conducted in vitro cleavage assays. Cleavage rates were measured for each aptazyme individually. Experiments were performed with and without theophylline to assess ligand-dependent activity. The results showed over 10 fold and 50 fold theophylline-induced cleavage increase after 1 hour incubation of *tetA* mRNA fragments of the TR/CA-GENA 1 and 2 respectively, with theophylline addition (**Figure 2**). This improvement highlights the effectiveness of the theophylline aptamer in activating ribozyme activity. Minimal cleavages were observed in the absence of theophylline with design TR/CA-GENA 1 and 2, consistent with prior reports of leakage activity in theophylline aptazymes. For example, Soukup et al. (2000) observed a potent activation of the hammerhead ribozyme—approximately 3000-fold—in the presence of theophylline, but also noted leakage rates of approximately 10_-_⁴ to 10_-_³ min_-_¹ in absence of theophylline. This leakage is attributed to a subpopulation of ribozymes that fold in an active conformation even without ligand binding, which can lead to low-level cleavage. No cleavage was detected for TR/CA-GENA 1 and Scr control designs.

**Figure 2:**
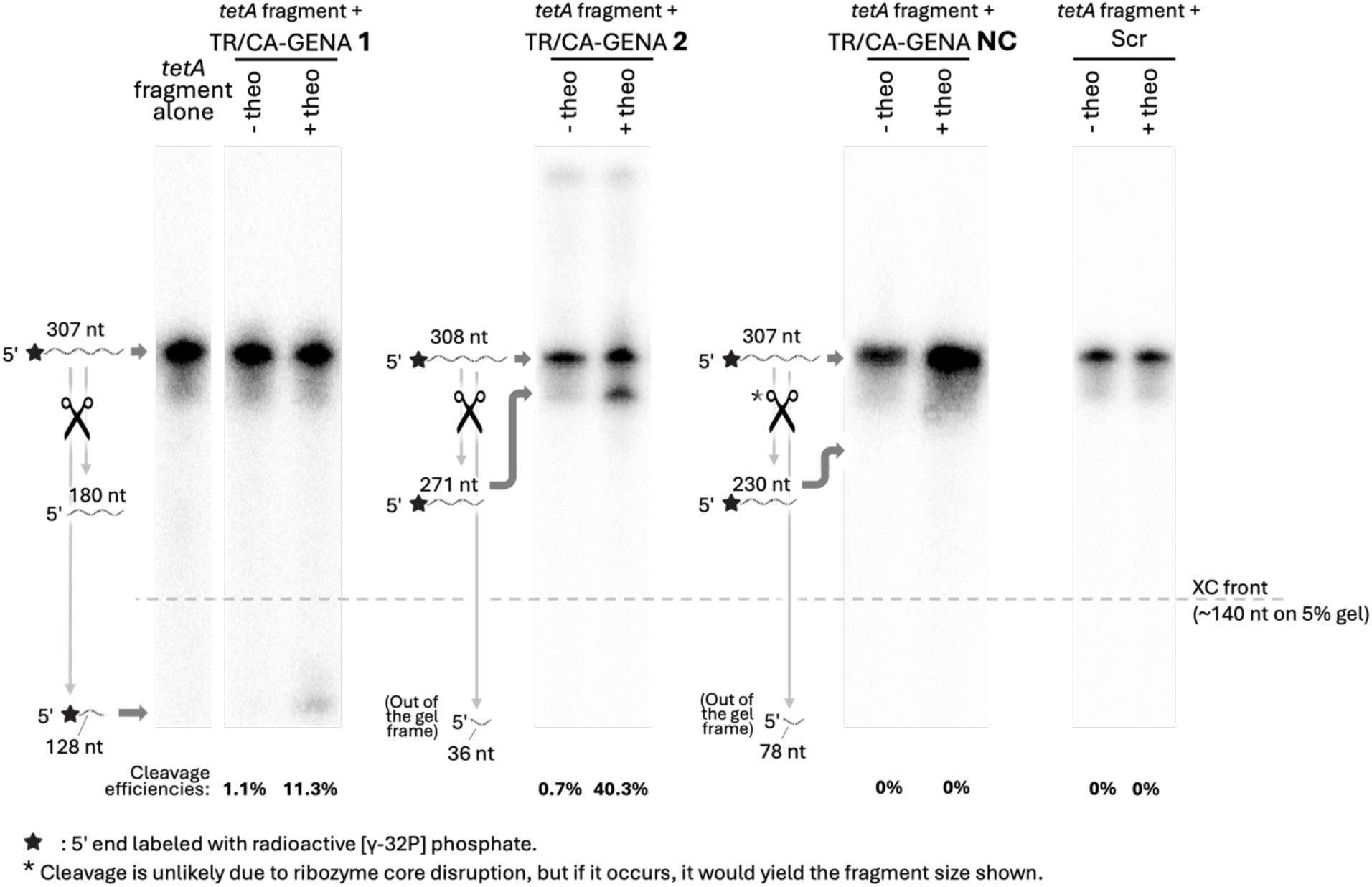
Aptazymes Cleave *tetA* RNA Fragments in a Trans-Acting Model. Autoradiogram of 5% denaturing PAGE showing cleavage of 5′ [γ-^32^P]-labeled *tetA* RNA fragments after 1-hour incubation with trans-acting aptazyme designs TR/CA-GENA-1, TR/CA-GENA-2, as well as controls TR/CA-GENA-NC (mutant HHR) and Scr (scrambled target sequence), with or without theophylline. Cleaved 5′ fragments appear as lower bands at expected positions, illustrated on the left of each gel. Cleavage efficiencies, calculated from the ratio of cleaved to uncleaved band intensities, are shown below each lane.

### Single Aptazyme Constructs Function as Effective GENAs in cells

To evaluate the in vivo efficacy of our aptazyme designs, we examined the impact of each construct placed in the 5′ UTR of the *tetA* gene mRNA (**Figure 1B**) on *Escherichia coli* strains DH5α and Top10. We measured bacterial growth rates under varying concentrations of tetracycline (Tc) and nickel (Ni²⁺). The goal was to determine whether the aptazymes, triggered by theophylline, could influence *tetA*-mediated phenotypes, including decreased Tc resistance and increased Ni²⁺ sensitivity like reported by Muranaka *et al.*^42^ (**Figure 3A**). Controls included no-GENA plasmids (constitutively expressing *tetA*) and no-plasmid clones, which lacked the *tetA* gene entirely. Experiments were conducted in absence and presence of theophylline to evaluate the ligand-dependent effects of our designs.

**Figure 3:**
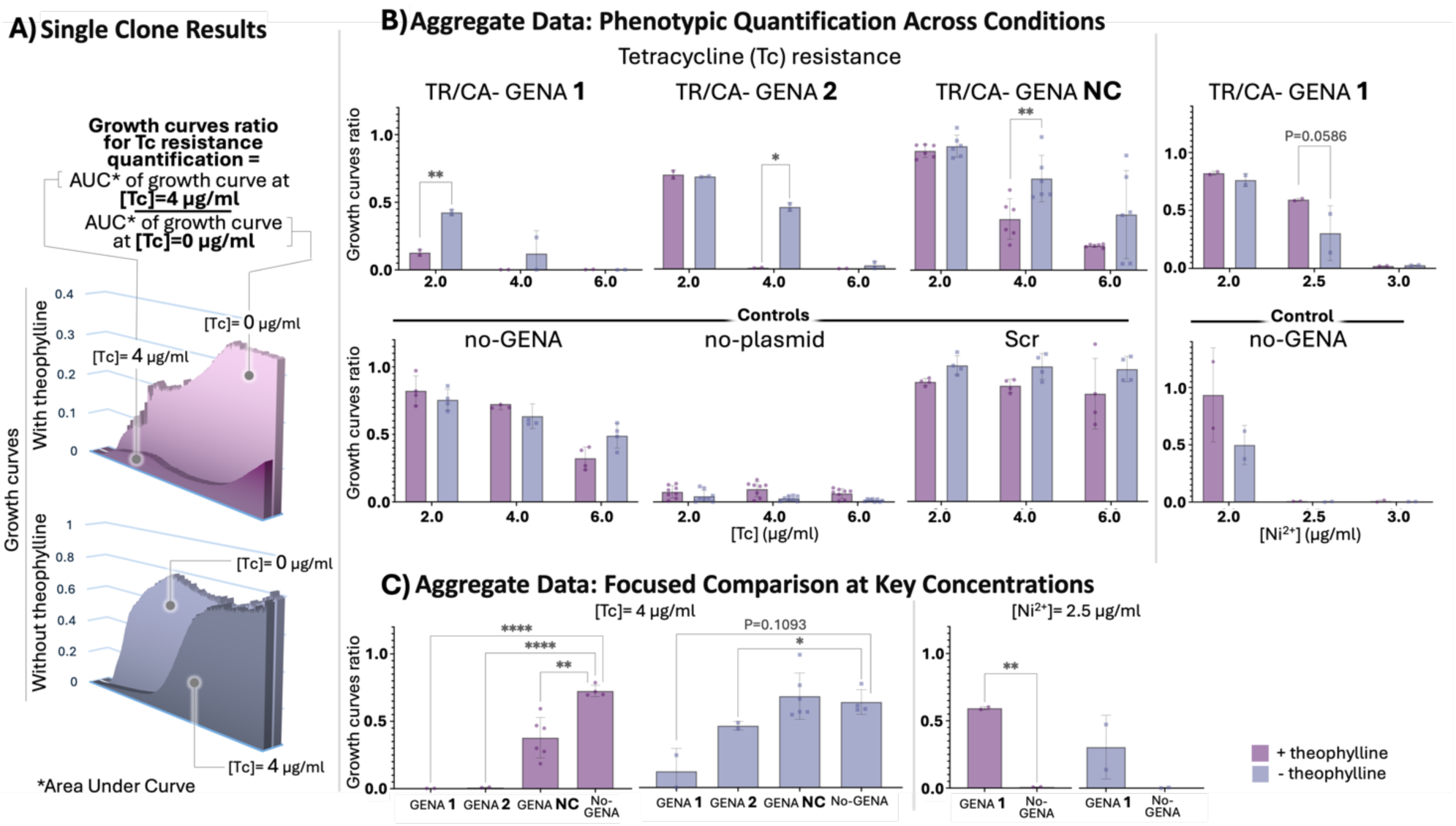
Aptazymes allow a theophylline-dependent control of growth under selective pressure from tetracycline or nickel. **A)** Growth curve analysis for single aptazyme clones: growth curves for a representative clone (TR/CA-GENA NC) are shown in the presence (lavender) and absence (blue/grey) of theophylline. Each graph presents two overlapping growth curves: one measured at Tc = 0 µg/mL (background curve) and another at Tc = 4 µg/mL (foreground curve). The method used to calculate the growth curve ratio, which reflects the phenotypic outcome (tetracycline resistance or nickel sensitivity), is detailed in the figure. **B)** Aggregated phenotypic data for tetracycline and nickel sensitivity: histograms display the mean growth curve ratios for three distinct GENAs (GENA NC, GENA 1 and GENA 2). **C)** This section focusses on conditions Tc 4 µg/ml and Ni²⁺ 2.5 µg/ml that showed the strongest phenotypic divergence. Test constructs (GENA 1, 2 and NC) are directly compared to the no-GENA control, allowing visual comparison and statistical testing of phenotypic significance under induced vs uninduced conditions. Each point on the histograms represents an individual replicate. Statistically significant differences were assessed using Welch’s t-test. Significance is indicated as follows: *p < 0.05, **p < 0.01. No plasmid controls are expected to be sensitive to Tc, no-GENA and scrambled aptazyme (Scr) are expected to be resistant (further discussion of this result is provided in Supplementary Note 1).

For quantitative analysis, we calculated the area under the growth curve (AUC) for each experimental condition and normalized it to the maximum growth observed under no-selection conditions (e.g., Tc = 0 μg/mL; **Figure 3A**). Specifically, for each tested condition, the AUC measured under selective pressure (e.g. with Tc or Ni) was divided by the AUC obtained under the corresponding non-selective condition. This generated a growth curve ratio that reflects how closely a clone’s growth under stress resembles its optimal growth, providing a standardized metric of phenotypic behavior—whether resistance or sensitivity. This approach provided a normalized growth rate metric, enabling a clear comparison across experimental conditions (**Figures 3B** and **C**), this normalization was notably necessary because theophylline has a mild impact on growth. This normalization method was particularly valuable for evaluating survival rates under selection pressures, an essential aspect of our study’s potential application in SELEX.

Results with the DH5alpha strain demonstrated a clear decrease in *tetA*-conferred Tc resistance for aptazyme designs compared to the no-GENA control plasmid (**Figure 3A** and **B**). In the presence of theophylline, all three designs showed a sharp decrease in average growth rates at Tc 4 μg/ml, ranging from quasi absence of growth up to approximately 50% of the growth observed in absence of theophylline. At Tc 4 μg/ml with theophylline, the growth rate of TR/CA-GENA 1 was reduced by more than 300-fold compared to no-GENA, while TR/CA-GENA 2 showed an 80-fold reduction (**Figure 3C**). In contrast, controls without aptazymes (no-GENA and scrambled) maintained growth rates above 80% under the same experimental conditions. The TR/CA-GENA 1 design demonstrated enhanced potency, exhibiting Tc sensitivity even at lower concentrations. At Tc 2 μg/ml in the presence of theophylline, growth for TR/CA-GENA 1 dropped below 15%. Notably, the TR/CA-GENA NC design, despite containing a disrupted ribozyme core and showing no detectable cleavage in vitro (**Figure 2**), also reduced growth under Tc selection, suggesting that it continues to interfere with *tetA* expression, possibly via minimal cleavage activity or by sterically hindering translation through binding at its target site. At Tc 4 μg/ml with theophylline, TR/CA-GENA NC reduced growth by 1.9-fold compared to the no-GENA control (**Figure 3C**). Similar results were obtained in a TOP10 *E. coli* strain (**Table S1, Figure S1** and **Supplementary Note 2**).

We also observed Tc sensitivity to varying degrees in assays without theophylline for all three aptazyme designs, manifesting at different Tc concentrations depending on the aptazyme-containing clones. In particular, TR/CA-GENA 1 growth rates were inhibited below 50% even without theophylline at a low Tc concentration of 2μg/ml. These results in the absence of theophylline indicate leakage activity, consistent with our in vitro findings (**Figure 2**), suggesting that a fraction of aptazymes can cleave their target even without the triggering ligand. However, since in vitro assays revealed only minimal cleavage activity in the absence of theophylline, we cannot exclude the possibility that the leakage-like effect observed with the TR/CA-GENA NC design is also partially driven by aptazyme binding alone. Given that aptazyme binding to the target gene could itself interfere with translation, it is plausible that such binding—though typically considered ligand-independent—might contribute to gene repression even in the absence of theophylline. Quantitatively, in the absence of theophylline, fold reductions in growth rate compared to the no-GENA control were 5.3-fold for TR/CA-GENA 1, 1.4-fold for TR/CA-GENA 2, and 0.9-fold (no-GENA growth rate lower) for TR/CA-GENA NC, indicating that even without the ligand, repression is present but much weaker (**Figure 3C**).

Despite the observed leakage activity in theophylline-free assays, the differences between theophylline-positive and theophylline-negative conditions remained significant, with higher growth rate inhibition observed in the presence of theophylline (for each aptazyme design, at least one tested Tc concentration yielded a statistically significant difference between theophylline-positive and theophylline-negative conditions in terms of growth rate reduction). The comparison between induced and uninduced conditions further highlights the impact of theophylline: TR/CA-GENA 1 and TR/CA-GENA 2 showed both a ∼50-fold increase in growth inhibition when comparing theophylline-positive to theophylline-negative conditions. In contrast, with TR/CA-GENA NC we observe only a 2-fold amplification, indicating that ligand-dependent switching greatly enhances repression efficiency for cleaving designs, but also for the inactive ribozyme core to a lesser extent. This theophylline-dependent effect was consistent across experiments, even though large standard deviations were sometimes noted due to experimental batch variations (within a single experimental batch, the comparison of data from the same clone in assays with and without theophylline showed more significant differences, see supplementary data, **Table S1** for raw data and **Figure S1** for single experimental batch growth curve comparisons). Interestingly, this trend also applied to the TR/CA-GENA NC design, which is not expected to cleave due to a disrupted ribozyme core. A plausible explanation is that the aptazyme’s binding to the target site interferes with translation, independently of its catalytic function. This interaction, even in the absence of cleavage, may still disturb ribosome access or mRNA structure sufficiently to reduce gene expression. The observed effect of theophylline may be due to an improved folding of the aptazyme, improving its binding even in absence of catalytic activity and thereby amplifying its regulatory impact. However, the magnitude of this effect remains limited: the 1.8-fold amplification observed for GENA NC is modest when compared to the ∼50-fold amplifications seen with TR/CA-GENA 1 and 2, respectively, reflecting the importance of cleavage activity in achieving strong repression.

To address potential alternative explanations for the observed gene regulation, we conducted additional control experiments. We included a scrambled control (Scr-control) based on the TR/CA-GENA 1 design but with a scrambled 5’ arm to prevent hybridization with the *tetA* sequence (**Figure S2**). This control exhibited a growth rate profile similar to the no-GENA control clone, supporting that specific aptazyme hybridization is essential for gene regulation. Furthermore, we explored the possibility that the observed effects might be due to non-specific interactions or structure effects in the 5’UTR region. To rule out this hypothesis, we performed experiments with *tetA* coding sequence-targeting ribozyme designs (**Table S1** and **Figure S1**), i.e. constitutively cleaving ribozymes. These ribozymes demonstrated a clear reduction in *tetA* expression in a fully theophylline-independent manner, reinforcing that gene regulation can occur through direct cleavage of the coding sequence by RNA tools that are not responsive to the ligand.

Taken together, our results not only demonstrate the efficacy of aptazymes for small-molecule-mediated gene regulation in a coding sequence-targeted, semi-trans acting configuration, but also—through the integration of in vitro cleavage data, in silico predictions, and in vivo phenotypic outcomes—provide insights into parameters that may predict aptazyme performance in cellular contexts (**Supplementary Table S2** and **Note 3**).

### TR/CA-GENA 1 Confers Theophylline Dependent Nickel Sensitivity

Nickel sensitivity assays with TR/CA-GENA 1 clones showed reduced sensitivity compared to no-GENA controls. At Ni²⁺ 2.5 μg/mL, TR/CA-GENA 1 clones demonstrated growth curve ratio of ∼60% with theophylline and ∼30% without, while no-GENA controls showed <5% growth under identical conditions. In the presence of theophylline, TR/CA-GENA 1 clones exhibited a 76-fold higher growth compared to no-GENA controls, a difference that was statistically significant (p < 0.01) (**Figure 3C**). Moreover, TR/CA-GENA 1 showed approximately 2-fold higher growth with theophylline compared to without, indicating that theophylline enhances repression of *tetA*, thereby mitigating the toxic effects of nickel stress. These results confirm the aptazyme′s regulatory effect on *tetA*, as increased cleavage reduces Ni²⁺ sensitivity (**Figure 3B**). Theophylline significantly improved growth rates in these assays, aligning with our expectation that theophylline triggers aptazyme-mediated cleavage of *tetA*, which is the source of nickel sensitivity. Although TOP10 is reported to be slightly more stress-tolerant than DH5α and may differ in Ni²⁺ sensitivity [44]^44^ we observed similar Ni²⁺-related effects in both strains (**Table S1, Figure S1** and **Supplementary Note 2**). Importantly, these results correlate with our observations in *E. coli* DH5α, further supporting the consistency of this regulatory mechanism. The reproducibility of these findings across different *E. coli* strains suggests that our gene regulation system is portable and functions effectively in multiple genetic backgrounds.

In brief, our in vivo tests demonstrated significant phenotypical effects of our aptazyme designs on both tetracycline resistance and nickel sensitivity. These effects were evident in the distinct survival profiles of TR/CA-GENA clones compared to control clones (no-GENA and no-plasmid) under comparable conditions, as well as in the differences observed between theophylline-positive and theophylline-negative assays for TR/CA-GENA clones.

### Multiple aptazyme construct assays

To further investigate the potential of our aptazyme designs, we constructed plasmids containing multiple aptazymes in the 5’UTR of the *tetA* gene. Two configurations were developed: a double aptazyme construct (TR/CA-GENA NC+2) incorporating both TR/CA-GENA NC and TR/CA-GENA 1, and a triple aptazyme construct (TR/CA-GENA NC+2+3) which added TR/CA-GENA 2 to the double construct (**Figure 4A**). In these constructs, we strategically positioned the aptazymes with spacers exceeding 20 nucleotides between each aptazyme (**Figure 4A**). Additionally, the distance between the aptazyme target cleavage sites was maintained at over 200 nucleotides. This design strategy aimed to minimize potential steric hindrance and enable simultaneous action of each aptazyme on its respective target site. For the double aptazyme construct, we positioned TR/CA-GENA NC and TR/CA-GENA 1 in a logical order from the 5’ end of the 5’UTR, allowing for a hairpin-like structure that facilitates proper alignment of the aptazymes with their target sites (**Figure 4A**). However, in the triple aptazyme construct, we deliberately placed TR/CA-GENA 2 in a position that did not follow this linear arrangement. This design choice was made to assess whether the spacing strategy we employed provides sufficient flexibility for the multiple aptazyme construct to properly bind their target sites and overcome potential positional constraints within the 5’UTR region, allowing all aptazymes to operate concurrently.

**Figure 4:**
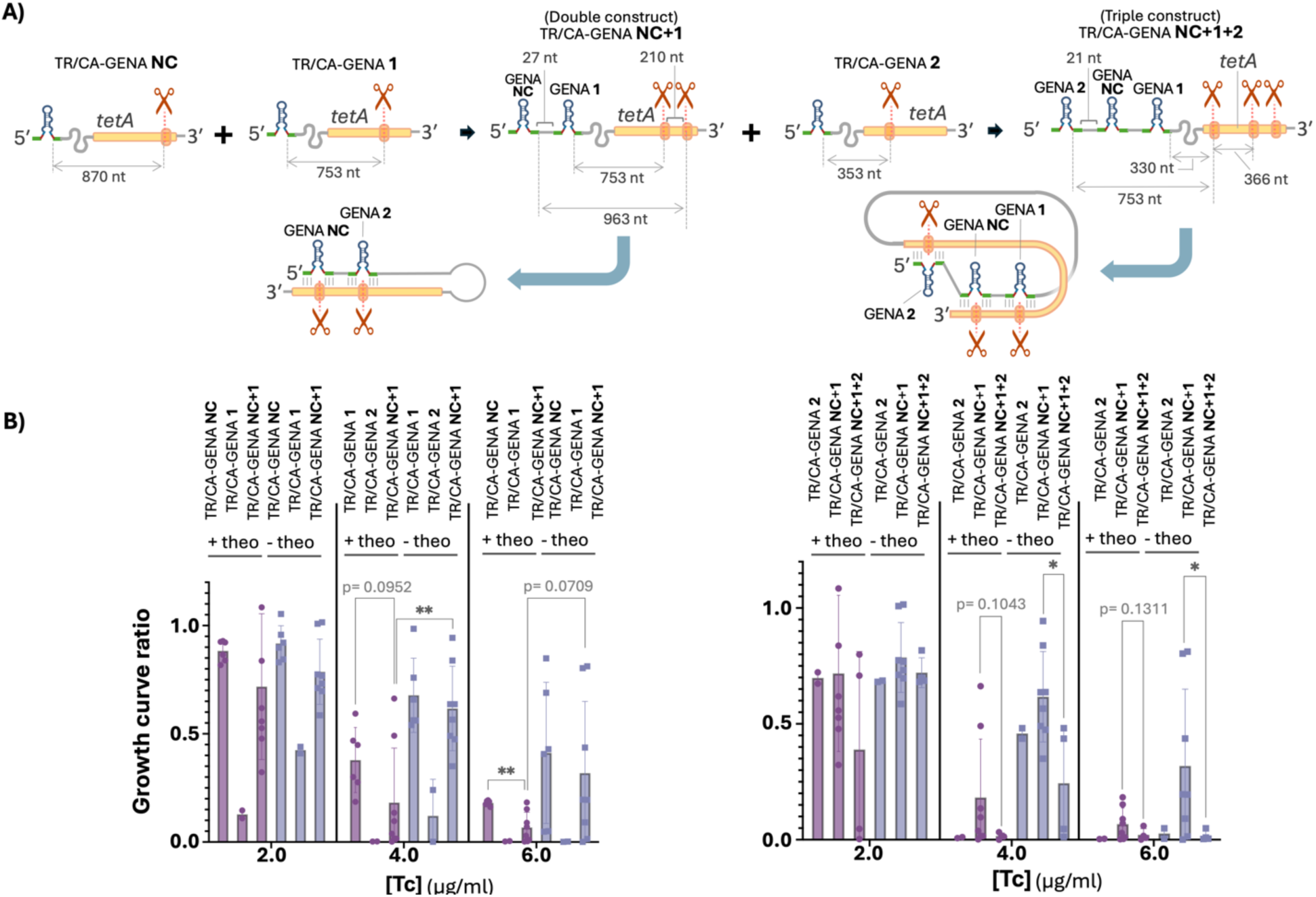
Design and Functional Characterization of Multiple Aptazyme Constructs. **A)** Construct design: Schematic representations of the different aptazyme configurations, showing how multiple aptazymes were combined within the 5’ UTR of *tetA*, including the distances between them and their respective cleavage sites. The left panel shows a construct where two aptazymes are arranged in a linear sequence with inter-aptazyme spacers, while the right panel depicts the triple aptazyme design in which aptazymes are arranged in a non-linear configuration. **(B)** Growth rates at increasing Tc concentrations for different constructs. To facilitate direct visual comparison, multiple aptazyme constructs are placed adjacent to the single aptazyme constructs from which they were derived. Statistical comparisons are indicated only when the multiple aptazyme constructs exhibit a reduction in Tc resistance, as expected. Differences are tested between the +Theo and -Theo conditions for each construct at the same [Tc], as well as between multiple aptazyme constructs and the single aptazyme constructs that served as their basis. Significance is denoted as follows: *p* < 0.05 (*), *p* < 0.01 (**), or exact *p*-value when 0.05 < *p* < 0.15 (Welch’s t-test, two-tailed).

The results of our experiments with multiple aptazyme constructs (**Figure 4B**) demonstrated significant effects on tetracycline (Tc) resistance in theophylline-positive assays. Both TR/CA-GENA NC+1 and TR/CA-GENA NC+1+2 constructs exhibited markedly reduced growth rates compared to the corresponding no-GENA control assays (**Figure 3**). For the TR/CA-GENA NC+1 construct, we observed a significant decrease in growth rates at Tc concentrations of 4 μg/ml and above, relative to the no-GENA control. The TR/CA-GENA NC+1+2 construct showed even greater sensitivity, with growth rates substantially affected at Tc concentrations as low as 2 μg/ml. Both constructs demonstrated responsiveness to theophylline, with growth rate inhibition more pronounced in theophylline-positive assays. However, it is worth noting that for the TR/CA-GENA NC+1+2 construct, the leaking cleavage activity resulted in similar growth rate outcomes for both theophylline-positive and theophylline-negative conditions, with growth rates approaching zero in both cases.

To assess the cumulative effect of multiple aptazymes, we arranged the growth rate data for multiple aptazyme constructs alongside their corresponding single aptazyme constructs (**Figure 4**). This is for a clear evaluation of the cumulative impacts of multiple aptazymes, for instance to gain insight into potential cooperative or additive effects of multiple aptazyme elements. Interestingly, the TR/CA-GENA NC+1 construct did not exhibit lower growth rates than TR/CA-GENA 1 alone. Instead, its performance appeared to be an average of TR/CA-GENA NC and TR/CA-GENA 1, with growth rates lower than TR/CA-GENA NC but higher than TR/CA-GENA 1. This suggests potential interference between the aptazymes in the double construct, diminishing the potent inhibitory effect observed with TR/CA-GENA 1 alone. Conversely, the TR/CA-GENA NC+1+2 construct, which combines the TR/CA-GENA 2 aptazyme with the TR/CA-GENA NC+1 double aptazyme construct, consistently demonstrated equal or stronger effects on *tetA* gene expression compared to TR/CA-GENA 2 alone and the double aptazyme construct TR/CA-GENA NC+1. At each tetracycline concentration, for both theophylline-positive and negative assays, TR/CA-GENA NC+1+2 exhibited the lowest growth rates. Although it did not reach the inhibition levels of TR/CA-GENA 1 alone, there was a clear pattern of improved performance compared to TR/CA-GENA NC+1 with the addition of TR/CA-GENA 2.

We hypothesize that in this triple aptazyme system, while one aptazyme may interfere with another, the third is momentarily free to operate. This scenario potentially allows the strongest aptazyme design, TR/CA-GENA 1, to be less inhibited than in the double aptazyme system, leading to the observed results. Conversely, observed results could be due to the cumulative effect of adding GENA 2 to the GENA NC+1 construct. These observations collectively suggest that combining aptazymes as we did does not produce a simple cumulative effect where each additional aptazyme linearly increases the overall regulatory impact. Instead, there could be interference between co-located aptazymes. This is not entirely surprising, given that the aptameric portions of the aptazymes are relatively highly structured sequences with base-pairing sections that may create inter-aptazyme interactions in multiple aptazyme constructs.

Nevertheless, the results indicate that aptazymes can function together in these multi-aptazyme systems. We suppose that each aptazyme in the construct may execute a portion of its original activity, with the TR/CA-GENA NC+1+2 results suggesting a partial restoration of TR/CA-GENA 1 activity. This may occur in a system where each aptazyme functions when not engaged in interference with others, implying that our design allows aptazymes to maintain functionality to some degree, even in configurations requiring complex folding like the triple aptazyme construct. These results suggest that optimization to minimize inter-aptazyme interference is required to accomplish better additive effect with multiple aptazymes. For systems employing aptazymes with aptameric parts responding to different ligands, we might expect reduced interference due to lower likelihood of base-pair matching between aptamers.

## Conclusion

Our study demonstrates the effectiveness of a semi-trans aptazyme-based gene regulation system, leveraging theophylline-responsive hammerhead ribozymes to cleave within coding sequences. Unlike conventional UTR-targeted approaches, this strategy enables portable and adaptable gene regulation, independent of promoter elements. By targeting the *tetA* gene in *E. coli*, we validated that ligand-dependent cleavage modulates gene expression, leading to predictable phenotypic changes in tetracycline resistance and nickel sensitivity.

The in vivo results confirm that aptazyme activity correlates with phenotypic effects, highlighting its potential for in vivo SELEX as a means to evolve new ligand-responsive regulatory elements. The distinct survival patterns observed under tetracycline and nickel selection pressures suggest that this system —building on the established use of the *tetA* gene in in vivo SELEX by Yokobayashi’s lab ^42^—could be directly applied in high-throughput selection strategies. By integrating aptazyme-mediated cleavage into in vivo SELEX, it would be possible to directly select aptazymes that regulate gene expression by targeting and cleaving coding sequences in a cellular context. Since this method would allow for direct functional selection, it could yield aptazymes that (i) respond to new ligands of interest and (ii) are inherently more effective, as they would be optimized through selective pressure rather than being derived from the rationally designed combination of a pre-existing aptamer and ribozyme. Unlike the model used in this study, where an existing aptamer was repurposed to function within an aptazyme, an in vivo SELEX approach could lead to the discovery of de novo aptazymes that are intrinsically better suited for ribozyme-mediated cleavage, potentially achieving higher efficiency and regulatory precision.

Our exploration of multi-aptazyme constructs revealed that while some designs showed interference effects, others demonstrated enhanced regulatory activity, particularly the triple aptazyme construct, which exhibited stronger inhibition than individual aptazymes. This study not only expands the toolkit for gene regulation but also highlights the untapped potential of coding sequence-targeted aptazymes, paving the way for more versatile and universal genetic control systems. By developing a portable and adaptable aptazyme system that can target multiple sites within coding sequences, we provide researchers with a powerful tool that is not limited to specific regions of the gene. This approach allows for greater flexibility in design and application, as it can be used to regulate gene expression by targeting various locations along the coding sequence, rather than being constrained to untranslated regions or specific regulatory elements. This versatility enhances the system’s potential for fine-tuned regulation and broadens its applicability across different genes and organisms. Further optimization of multi-aptazyme interactions, ligand compatibility, and spatial arrangement will likely enhance their effectiveness in synthetic biology and gene therapy. By demonstrating that coding sequences can serve as efficient regulatory targets, this study lays the groundwork for more flexible, portable, and tunable gene control systems, expanding the potential of nucleic acid-based gene regulation across diverse applications.

## Materials and Methods

### Plasmid Construction and Cloning

The plasmid pLac-thiM-ON-tetA-gfpuv (plasmid map in **Figure S3**), kindly provided by the Prof. Yohei Yokobayashi (Okinawa Institute of Science and Technology), was used as the backbone for this study. This vector contains a mutated riboswitch (derived from the *E. coli thiM* thiamine pyrophosphate riboswitch) that is always in the ON conformation, which drives the expression of a fused *tetA*–GFPuv gene ^42,45^. The “ON” designation indicates that this riboswitch mutant never blocks expression, leading to continuous *tetA*-GFPuv expression. Aptazyme designs were generated using Ribosoft software ^32^, with manual elongation of binding arms to enhance hybridization strength to the *tetA* target gene. Inserts were obtained by PCR amplification using primers and templates listed in **Table S3**. These PCR products were designed with EcoR1, Pst1 and/or Xma1 restriction sites, enabling directed cloning into the target plasmid. To obtain the multiple aptazyme constructs, we employed a stepwise cloning strategy that allowed the successive integration of aptazyme designs into the plasmid backbone. First, the TR/CA-GENA 1 insert was introduced into the plasmid already containing TR/CA-GENA NC using the XmaI restriction site, generating the TR/CA-GENA NC+1 construct. This intermediate plasmid was then used as the backbone for the final construct, using the PstI restriction site to incorporate TR/CA-GENA 2, yielding the TR/CA-GENA NC+1+2 construct. This sequential approach ensured precise assembly of multiple aptazyme designs within the same plasmid. Cloning was performed using homemade chemically competent *E. coli* DH5α cells, prepared using the standard calcium chloride protocol or commercial TOP10 (Thermo Fisher). Transformation was performed by heat shock, and recombinant clones were selected on LB-agar plates supplemented with ampicillin (100 µg/mL). Successful cloning was verified by colony PCR and plasmid sequencing (Plasmidsaurus Inc.). The TR/CA-GENA NC construct, which contains a disrupted ribozyme core due to an extra guanine insertion, was not deliberately designed. This mutation was discovered during sequencing analysis of one of the clones, and we chose to retain it in the study as a useful non-cleaving control to assess the specific contribution of ribozyme cleavage activity to gene regulation. No special mutagenesis or cloning steps were required to generate this variant.

### In Vitro Cleavage Assays

To generate RNA transcripts for in vitro cleavage experiments, PCR amplification was performed using primers designed to introduce a T7 promoter sequence upstream of the TR/CA-GENA constructs and control sequences (see **Table S3** for primer details). In vitro transcription was carried out using T7 RNA polymerase, provided by Sherbrooke University, following their standard protocol. The transcribed RNAs were purified by 6% denaturing polyacrylamide gel electrophoresis (PAGE, 8 M urea). Purified RNA probes were then 5′-end radiolabeled with γ-³²P ATP (Revvity) using T4 polynucleotide kinase (T4 PNK, New England Biolabs), following the manufacturer′s protocol.

For cleavage reactions, TR/CA-GENA and control design RNAs were mixed with target RNA in 50 mM Tris-HCl (pH 7.5 at 23°C), 20 mM MgCl₂ buffer. Samples underwent a heat shock treatment at 65°C for 4 minutes, followed by incubation at room temperature for durations specified in the Results section. Reactions were analyzed using 8% denaturing PAGE. Gels were exposed to phosphor imaging screens for 30 minutes and scanned with a Typhoon™ FLA9500 imaging system (GE Healthcare Life Sciences). Cleavage efficiency was quantified by measuring band intensities using ImageQuant TL software (GE Healthcare Life Sciences). The cleavage rate was determined by calculating the ratio of cleaved fragment intensity to the total band (cleaved and full length) intensity.

### In Vivo Growth Assays and Data Analysis

Growth assays were conducted using overnight cultures of *E. coli* DH5α or Top10 strains in LB broth supplemented with ampicillin (100 µg/mL). Overnight cultures were refreshed by diluting 1:100 in fresh LB medium and incubated at 37°C until reaching OD_600_ ∼1.0. The optical density at 600 nm (OD_600_) was measured using an Eppendorf BioSpectrometer®. To ensure consistent starting cell densities across conditions, cultures were normalized as follows: 1 mL of culture was prepared for each clone by taking 100 µL of an OD_600_ ∼1 culture, and other cultures were adjusted accordingly (e.g., if a culture had OD_600_ = 0.9, 111 µL was used and adjusted to 1 mL with LB). For growth assays, 5 µL of the normalized culture was transferred to 96-well plates, containing LB-ampicillin (except for no-plasmid controls) and supplemented with either 10 mM theophylline for DH5α or 8 mM theophylline for Top10, as well as varying concentrations of tetracycline (Tc) or nickel (Ni), as described in the Results section. Plates were incubated at 37°C in a Cytation 3 plate reader (BioTek), with 20 minute cycles of shaking and OD_600_ readings for 16–30 hours (depending on the experimental batch; see Supplementary Excel file for details).

To account for initial culture density when theophylline was added, the first recorded OD_600_ value of each sample was subtracted from subsequent OD_600_ values at each time point:

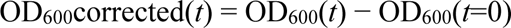

Where OD_600_(*t*=0) is the first OD_600_ reading taken for each sample.

To quantify bacterial growth over time, the area under the growth curve (AUC) was calculated using the trapezoidal rule. This method estimates AUC by summing the areas of small trapezoidal sections formed between consecutive OD_600_ time points. Trapezoidal AUC estimation is widely used in biological time-course analyses ^46^ and has been applied in bacterial growth monitoring contexts ^47^.The AUC formula used was:

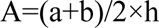

where:

- a = OD_600_ at time point *t*₁,
- b = OD_600_ at time point *t*₂,
- h = time interval (*t*₂ - *t*₁).

The AUC was determined by summing the area of each trapezoid between consecutive time points throughout the entire growth experiment.

Fold change calculations were based on the average growth curve ratios obtained from biological replicates. Growth curve ratios were calculated as described in Figure 3A, representing the area under the curve (AUC) normalized to the no-antibiotic control. For comparisons between different clones under the same experimental condition (e.g., with theophylline), fold changes were computed by dividing the mean growth ratio of one clone by that of the reference (e.g., no-GENA). For comparisons between theophylline-positive and -negative conditions for the same clone, fold changes were calculated by dividing the mean growth ratio with theophylline by the mean without.

## Data availability statement

The original contributions presented in this study are included in the article and its supplementary material; further inquiries can be directed to the corresponding author.

## Authors contributions

JP conceived the original idea and supervised the study. EY conceptualized the study and designed the experiments in discussion with JP. EY carried out most of the experiments and was responsible for troubleshooting technical issues. YN, FPD, and NES contributed to the experimental work. EY wrote the manuscript.

## Acknowledgments

We sincerely thank Yohei Yokobayashi for his generous gift of the plasmid pLac-thiM-ON-tetA-gfpuv, which was instrumental in the development of this study. The authors utilized ChatGPT (OpenAI, San Francisco, CA) to assist in refining the language and grammar of the manuscript.

All content was reviewed and edited by the authors to ensure accuracy and adherence to scientific standards.

