## Supporting information for "Harnessing Coding Sequence Cleavage: Theophylline Aptazymes as Portable Gene Regulators in Bacteria"

### Supplementary Note 1. Additional interpretation of Figure 3

To better understand the variability observed in Figure 3, we provide here further interpretation regarding the sources of variation between experimental replicates. The high standard deviations observed when combining experiments from different dates can be attributed to several factors: i) use of bacterial cultures potentially affected by passages ; ii) fluctuations in plasmid content in cultures; iii) Imprecise normalization of bacterial quantities across experiments due to limitations in OD measurement, especially considering exponential growth. These factors likely resulted in slight profile variations between experiments, leading to the observed standard deviations when data from different days were combined.

color scale: 1 / 0.8 / 0.6 / 0.4 / 0.2 / 0

### DH5a Tc resistance tests

#### Single aptzyme constructs

**TR/CA-GENA NC**

| | [Tc], ug/ml | 0 | 2 | 3 | 4 | 5 | 6 |
|---|---|---|---|---|---|---|---|
| 21-Jul-23 | + Theo | 1 | 0.86527629 | 0.49206904 | 0.27620231 | 0.19853323 | 0.18896783 |
| | + Theo | 1 | 0.81762284 | 0.46989767 | 0.18564685 | 0.16931515 | 0.18671815 |
| | - Theo | 1 | 0.9212774 | 0.73671274 | 0.56128065 | 0.33751701 | 0.0501944 |
| | - Theo | 1 | 0.95775103 | 0.7842701 | 0.54060775 | 0.34937507 | 0.04755762 |
| 28-Jul-23 | + Theo | 1 | 0.92190294 | 0.84321856 | 0.5930257 | 0.41167292 | 0.17448017 |
| | + Theo | 1 | 0.9247458 | 0.84851471 | 0.44855209 | 0.43833301 | 0.19311763 |
| | - Theo | 1 | 0.83023151 | 0.84948715 | 0.75928231 | 0.72534919 | 0.68948193 |
| | - Theo | 1 | 1.05365087 | 1.14910309 | 0.98506509 | 0.85683031 | 0.84859691 |
| 14-Jul-23 | + Theo | 1 | 0.83921651 | 0.64818334 | 0.30022359 | 0.20431805 | 0.16166387 |
| | + Theo | 1 | 0.92878976 | 0.7381336 | 0.46849562 | 0.27219857 | 0.17959645 |
| | - Theo | 1 | 0.8435725 | 0.89482387 | 0.56296479 | 0.37548376 | 0.40656234 |
| | - Theo | 1 | 0.89773524 | 0.89518282 | 0.65815373 | 0.22008498 | 0.42867531 |

**TR/CA-GENA 1**

| | [Tc], ug/ml | 0 | 2 | 4 | 6 |
|---|---|---|---|---|---|
| 12 janv 2024 | + Theo | 1 | 0.10953367 | 0.0025516 | 0.00433064 |
| | + Theo | 1 | 0.14614864 | 0.00218829 | 0.00267937 |
| | - Theo | 1 | 0.4396069 | 0.24027334 | 0.00086029 |
| | - Theo | 1 | 0.40926321 | 0.00132229 | 0.00194534 |

**TR/CA-GENA 2**

| | [Tc], ug/ml | 0 | 2 | 4 | 6 |
|---|---|---|---|---|---|
| 12 janv 2024 | + Theo | 1 | 0.67376011 | 0.00710783 | 0.00385256 |
| | + Theo | 1 | 0.72267491 | 0.01096784 | 0.00271856 |
| | - Theo | 1 | 0.68657985 | 0.43615689 | 0.00439575 |
| | - Theo | 1 | 0.68107998 | 0.48124105 | 0.0491808 |

#### Multiple aptzymes constructs

**GENA NC+1**

| | [Tc], ug/ml | 0 | 2 | 3 | 4 | 5 | 6 |
|---|---|---|---|---|---|---|---|
| 21-Jul-23 | + Theo | 1 | 0.55764373 | 0.3642442 | 0.09660112 | 0.07492119 | 0.07032359 |
| | + Theo | 1 | 0.83729196 | 0.43686102 | 0.1349079 | 0.11727041 | 0.08909264 |
| | - Theo | 1 | 0.71145899 | 0.67235031 | 0.47478142 | 0.19961669 | 0.01390619 |
| | - Theo | 1 | 0.76842157 | 0.71862398 | 0.49049048 | 0.34775399 | 0.09074667 |
| 28-Jul-23 | + Theo | 1 | 1.31806684 | 1.09436903 | 0.66291684 | 0.39086326 | 0.18294855 |
| | + Theo | 1 | 1.08421853 | 0.80848755 | 0.49083554 | 0.30203958 | 0.15217504 |
| | - Theo | 1 | 1.01514961 | 0.98615041 | 0.9436353 | 0.84980825 | 0.81173055 |
| | - Theo | 1 | 1.00837823 | 0.91091672 | 0.83849553 | 0.82345001 | 0.80321425 |
| 09-Feb-24 | + Theo | 1 | 0.61864644 | | 0.03807974 | | 0.02905473 |
| | + Theo | 1 | 0.5262461 | | 0.01697521 | | 0.0098805 |
| | - Theo | 1 | 0.7744573 | | 0.63660755 | | 0.19492665 |
| | - Theo | 1 | 0.73156128 | | 0.56575807 | | 0.19582756 |
| 12 janv 2024 | + Theo | 1 | 0.47783186 | | 0.00176308 | | 0.00202236 |
| | + Theo | 1 | 0.32244569 | | 0.01543677 | | 0.0035032 |
| | - Theo | 1 | 0.69706781 | | 0.63695983 | | 0.43664783 |
| | - Theo | 1 | 0.58570317 | | 0.34989753 | | 0.00063559 |

**TR/CA-GENA NC+1+2**

| | [Tc], ug/ml | 0 | 2 | 4 | 6 |
|---|---|---|---|---|---|
| 09-Feb-24 | + Theo | 1 | 0.0469854 | 0.01336552 | 0.05946773 |
| | + Theo | 1 | 0.00274529 | 0 | 0.00067011 |
| | - Theo | 1 | 0.57985969 | 0.00279959 | 0.00255672 |
| | - Theo | 1 | 0.49951343 | 0.06914935 | 0.00073815 |
| 12 janv 2024 | + Theo | 1 | 0.79929917 | 0.03238561 | 0.01660765 |
| | + Theo | 1 | 0.70805047 | 0.01748384 | 0.00822108 |
| | - Theo | 1 | 0.69904708 | 0.01201492 | 0.00294907 |
| | - Theo | 1 | 0.81678101 | 0.04617587 | 0.00908981 |

#### Controls

**no-plasmid****

| | [Tc], ug/ml | 0 | 2 | 3 | 4 | 5 | 6 |
|---|---|---|---|---|---|---|---|
| 21-Jul-23 | + Theo | 1 | 0.13543598 | 0.17808565 | 0.16452826 | 0.11018885 | 0.07920787 |
| | + Theo | 1 | 0.08821302 | 0.08109157 | 0.1159839 | 0.11222271 | 0.079526 |
| | - Theo | 1 | 0.02567171 | 0.03024298 | 0.02801988 | 0.01977124 | 0.0217209 |
| | - Theo | 1 | 0.02147369 | 0.02688796 | 0.0152495 | 0.01798682 | 0.00461233 |
| 28-Jul-23 | + Theo | 1 | 0.05358164 | 0.10777098 | 0.12382341 | 0.10678065 | 0.09143495 |
| | + Theo | 1 | 0.05985704 | 0.10175697 | 0.06601163 | 0.08834998 | 0.07158504 |
| | - Theo | 1 | 0.115029 | 0.05433967 | 0.04797517 | 0.02757706 | 0.02141306 |
| | - Theo | 1 | 0.1022968 | 0.04527361 | 0.04394449 | 0.03666374 | 0.00933094 |
| 14-Jul-23 | + Theo | 1 | 0.10087588 | 0.12992075 | 0.13196474 | 0.0997952 | 0.05671966 |
| | + Theo | 1 | 0.12744918 | 0.16686299 | 0.12428831 | 0.13576692 | 0.10412767 |
| | - Theo | 1 | 0.04650838 | 0.03768741 | 0.04206776 | 0.03229936 | 0.02036851 |
| | - Theo | 1 | 0.02524588 | 0.02446247 | 0.01843694 | 0.02387939 | 0.01516342 |
| 12 janv 2024 | + Theo | 1 | 0.00804387 | | 0.00822545 | | 0.00502969 |
| | + Theo | 1 | 0.02639502 | | 0.02704737 | | 0.01053794 |
| | - Theo | 1 | 0.00086869 | | 0.00469712 | | 0.00047707 |
| | - Theo | 1 | 0.00050659 | | 0.00335881 | | 0.00037036 |

**no-GENA***

| | [Tc], ug/ml | 0 | 2 | 3 | 4 | 5 | 6 |
|---|---|---|---|---|---|---|---|
| 21-Jul-23 | + Theo | 1 | 0.97127104 | 0.76749741 | 0.69533423 | 0.47501335 | 0.23719968 |
| | + Theo | 1 | 0.82059894 | 0.71842676 | 0.69876125 | 0.44031804 | 0.2679021 |
| | - Theo | 1 | 0.85916633 | 0.75270229 | 0.60805875 | 0.49437868 | 0.48511216 |
| | - Theo | 1 | 0.76449371 | 0.68936465 | 0.57683929 | 0.49847242 | 0.36615683 |
| 28-Jul-23 | + Theo | 1 | 0.70835764 | 0.85823031 | 0.78677594 | 0.39890726 | 0.40601009 |
| | + Theo | 1 | 0.79069032 | 0.66711868 | 0.72036595 | 0.46895567 | 0.38240399 |
| | - Theo | 1 | 0.67154206 | 0.65337494 | 0.58356428 | 0.5724522 | 0.52492535 |
| | - Theo | 1 | 0.72857548 | 0.60035883 | 0.77022464 | 0.67424043 | 0.58398049 |

**Scr*****

| | [Tc], ug/ml | 0 | 2 | 4 | 6 |
|---|---|---|---|---|---|
| 24mars 2024 | - theo | 1 | 0.98060157 | 0.95708995 | 0.89796178 |
| | - theo | 1 | 1.10305132 | 1.06168893 | 1.07041261 |
| | + theo | 1 | 0.84888188 | 0.89882592 | 0.65837502 |
| | + theo | 1 | 0.88272948 | 0.79705906 | 0.5690956 |
| | - theo | 1 | 0.93386276 | 0.88245459 | 0.89496987 |
| | - theo | 1 | 1.00346733 | 1.08867076 | 1.04774194 |
| | + theo | 1 | 0.91181301 | 0.83409433 | 0.79520363 |
| | + theo | 1 | 0.88841976 | 0.89039324 | 1.16171927 |

**Rbz******

Ratio compared to [Tc]=0 values

| | [Tc], ug/ml | 0 | 2.1 | 2.8 | 3.75 | 5 |
|---|---|---|---|---|---|---|
| 24mars 2024 | Rbz - theo | 1 | 0.95152038 | 0.09596445 | 0.01770733 | 0 |
| | Rbz - theo | 1 | 0.99439568 | 0.0729237 | 0.01242404 | 0 |
| | Rbz + theo | 1 | 1.0235732 | 0.62853598 | 0.10967742 | 0 |
| | Rbz + theo | 1 | 1.96893939 | 0.9875 | 0.15340909 | 0 |

* no-GENA: clones transformed with no aptazyme containing plasmid (expressing tetA constitutively)

**no-plasmid: no-transformed bacteria (no tetA gene)

***Scr: scrambled aptazyme.

### Top10 Tc resistance tests

#### Single aptzyme constructs

**TR/CA-GENA NC**

| | [Tc], ug/ml | 0 | 0.6 | 1.2 | 2.4 | 4.8 |
|---|---|---|---|---|---|---|
| 26-Jan-22 | clone2 +theo | 1 | 1.04853781 | 1.03599239 | 0.55675722 | 0.07397474 |
| | clone2 -theo | 1 | 0.98809524 | 1.00188895 | 0.65616763 | 0.17795642 |

| | [Tc], ug/ml | 0 | 2 | 2.5 | 3 | 3.5 | 4 |
|---|---|---|---|---|---|---|---|
| 11Fev2022 | + Theo | 1 | 0.40035457 | 0.18144027 | 0.0927375 | 0.0469495 | 0.02121286 |
| | - Theo | 1 | 0.54130596 | 0.22205524 | 0.14255477 | 0.02713515 | 0.0157477 |

#### Multiple aptzymes constructs

**TR/CA-GENA NC+1**

| [Tc], ug/ml | 0 | 2 | 2.5 | 3 | 3.5 | 4 |
|---|---|---|---|---|---|---|
| + Theo | 1 | 0.35146882 | 0.18798764 | 0.03424622 | 0.05168351 | 0.02173116 |
| - Theo | 1 | 0.57027401 | 0.44548117 | 0.24677064 | 0.03126064 | 0.01734432 |

#### Controls

**no-GENA***

| | [Tc], ug/ml | 0 | 0.6 | 1.2 | 2.4 | 4.8 |
|---|---|---|---|---|---|---|
| 26-Jan-22 | + Theo | 1 | 1.34313617 | 1.33769837 | 1.17234491 | 0.58450782 |
| | - Theo | 1 | 0.99721775 | 0.90457766 | 0.92931357 | 0.44059471 |

| | [Tc], ug/ml | 0 | 2 | 2.5 | 3 | 3.5 | 4 |
|---|---|---|---|---|---|---|---|
| 11Fev2022 | + Theo | 1 | 0.60361466 | 0.51458694 | 0.51988621 | 0.42232387 | 0.22117476 |
| | - Theo | 1 | 1.16543135 | 1.02545098 | 0.80783159 | 0.51798829 | 0.11188953 |

**no-plasmid****

| | [Tc], ug/ml | 0 | 0.6 | 1.2 | 2.4 | 4.8 |
|---|---|---|---|---|---|---|
| 26-Jan-22 | + Theo | 1 | 0.23289365 | 0.05873866 | 0.05173124 | 0.02061006 |
| | - Theo | 1 | 0.08067099 | 0.02718392 | 0.0197774 | 0.01088958 |

| | [Tc], ug/ml | 0 | 2 | 2.5 | 3 | 3.5 | 4 |
|---|---|---|---|---|---|---|---|
| 11Fev2022 | + Theo | 1 | 0.0077821 | 0.02613314 | 0.01908718 | 0.00515301 | 0.01330319 |
| | - Theo | 1 | 0.00678306 | 0.00518505 | 0.00472604 | 0.00385903 | 0.00703806 |

### Top10 Nickel sensitivity tests

#### Single aptzyme constructs

**TR/CA-GENA NC**

| | $[Ni^{2+}]$, ug/ml | 0 | 0.3 | 0.4 | 0.5 | 1 | 2 |
|---|---|---|---|---|---|---|---|
| 11Fev2022 | + Theo | 1 | 0.80284627 | 0.86847353 | 0.91914431 | 1.07151922 | 0.05556563 |
| | - Theo | 1 | 1.50363466 | 1.61749676 | 1.63304304 | 0.75478624 | 0.00834893 |

**TR/CA-GENA NC+1**

| | | | | | | |
|---|---|---|---|---|---|---|
| + Theo | 1 | 1.26180666 | 1.11590069 | 1.40369783 | 1.19830956 | 0.04828315 |
| - Theo | 1 | 1.47647234 | 1.40927879 | 1.1681492 | 0.72887801 | 0.00505936 |

#### Controls

**no-GENA***

| | | | | | | |
|---|---|---|---|---|---|---|
| + Theo | 1 | 1.28950485 | 1.26356302 | 1.50710532 | 0.69006735 | 0.02124195 |
| - Theo | 1 | 1.2595233 | 1.2403794 | 1.46692069 | 0.59875652 | 0.01932827 |

**no-plasmid****

| | | | | | | |
|---|---|---|---|---|---|---|
| + Theo | 1 | 1.49887059 | 1.19299212 | 0.42592695 | 0.76293317 | 0.01008209 |
| - Theo | 1 | 0.97833676 | 1.03166547 | 1.03505731 | 0.61201753 | 0.00982892 |

### DH5a Nickel sensitivity tests

**TR/CA-GENA 1**

| | $[Ni^{2+}]$, ug/ml | 0 | 2 | 2.5 | 3 |
|---|---|---|---|---|---|
| 21mars2024 | + Theo | 1 | 0.80977647 | 0.58476968 | 0.01981036 |
| | + Theo | 1 | 0.83388795 | 0.60210442 | 0.02206755 |
| | - Theo | 1 | 0.72205707 | 0.47061814 | 0.02638967 |
| | - Theo | 1 | 0.8029367 | 0.13665877 | 0.02888521 |

**no-GENA***

| | | | | |
|---|---|---|---|---|
| + Theo | 1 | 1.22526573 | 0.00865196 | 0.01613269 |
| + Theo | 1 | 0.6447792 | 0.00694104 | 1.3059E-05 |
| - Theo | 1 | 0.37671198 | 8.538E-05 | 0.00300586 |
| - Theo | 1 | 0.61899935 | 0.00501628 | 0.00411683 |

**Table S1. Comprehensive Growth Rate Data from All Experimental Conditions:** Complete dataset of growth rates obtained across all tetracycline (Tc) resistance and nickel sensitivity assays performed in this study. The data are organized by bacterial strain (DH5α or Top10), aptazyme construct type (single or multiple), and experimental conditions (+/- theophylline and varying Tc or $Ni^{2+}$ concentrations). The heatmap highlights growth rate variations, with higher growth rates in lighter shades and lower growth rates in darker red shades, allowing for rapid visual interpretation of tetracycline resistance or nickel sensitivity trends across conditions. Each experiment is labeled with its corresponding date and replicate, ensuring traceability of experimental results.

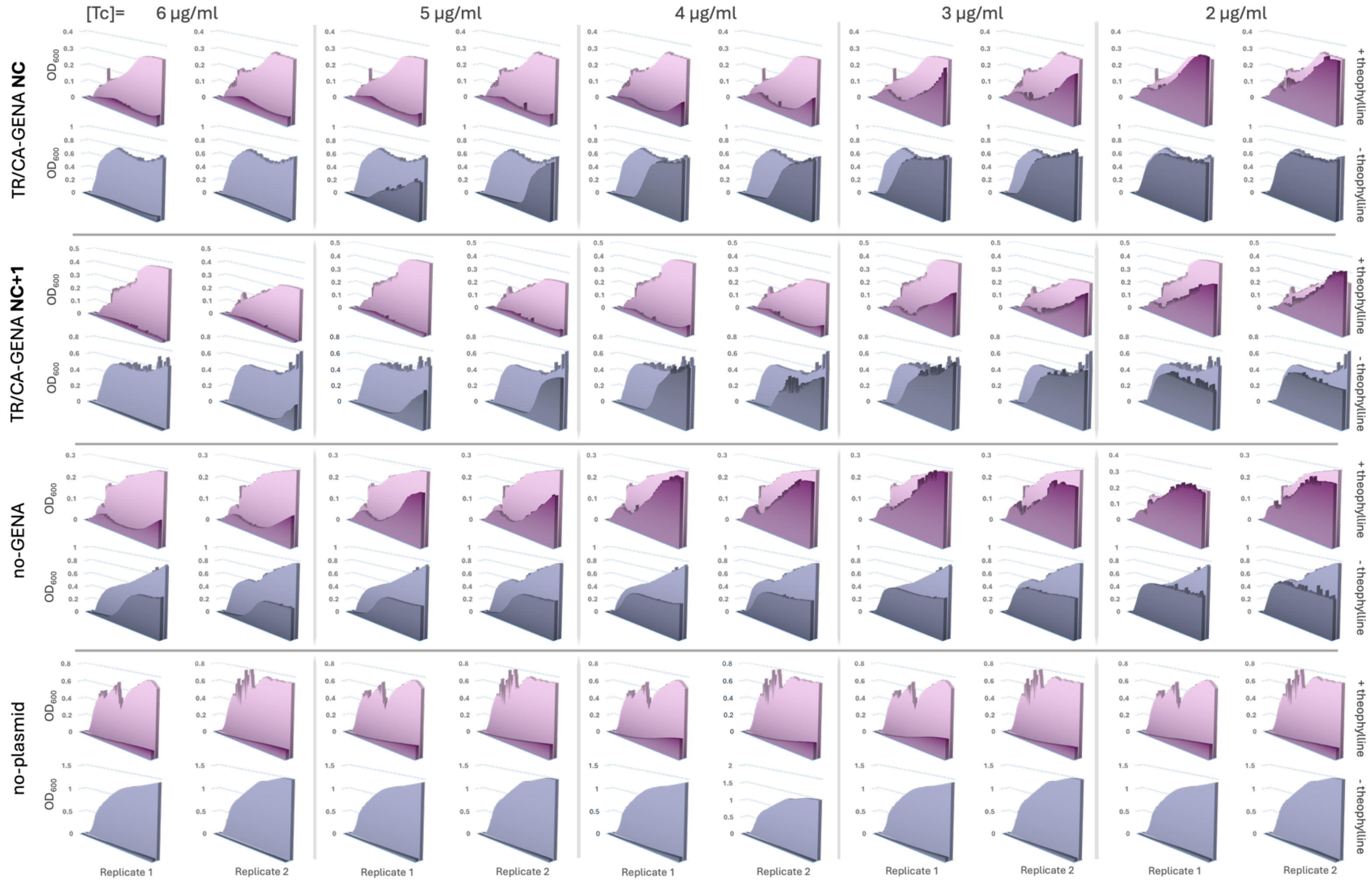

**Figure S1. Growth Curve Analysis of Tetracycline Resistance for Different Clones (part 1):** Growth curves for each clone used in this study are presented, showing tetracycline (Tc) resistance tests across different Tc concentrations. Each clone was tested in two replicates, with growth recorded in + theophylline (lavender curves) and − theophylline (blue-grey curves) conditions. To enable direct comparison, each growth curve is displayed alongside its corresponding [Tc] = 0 µg/mL reference curve (background curve), representing the same clone under identical experimental batch conditions. This reference allows for assessing the impact of Tc stress relative to the normal growth baseline. The figure is split across two pages to accommodate all clones tested.

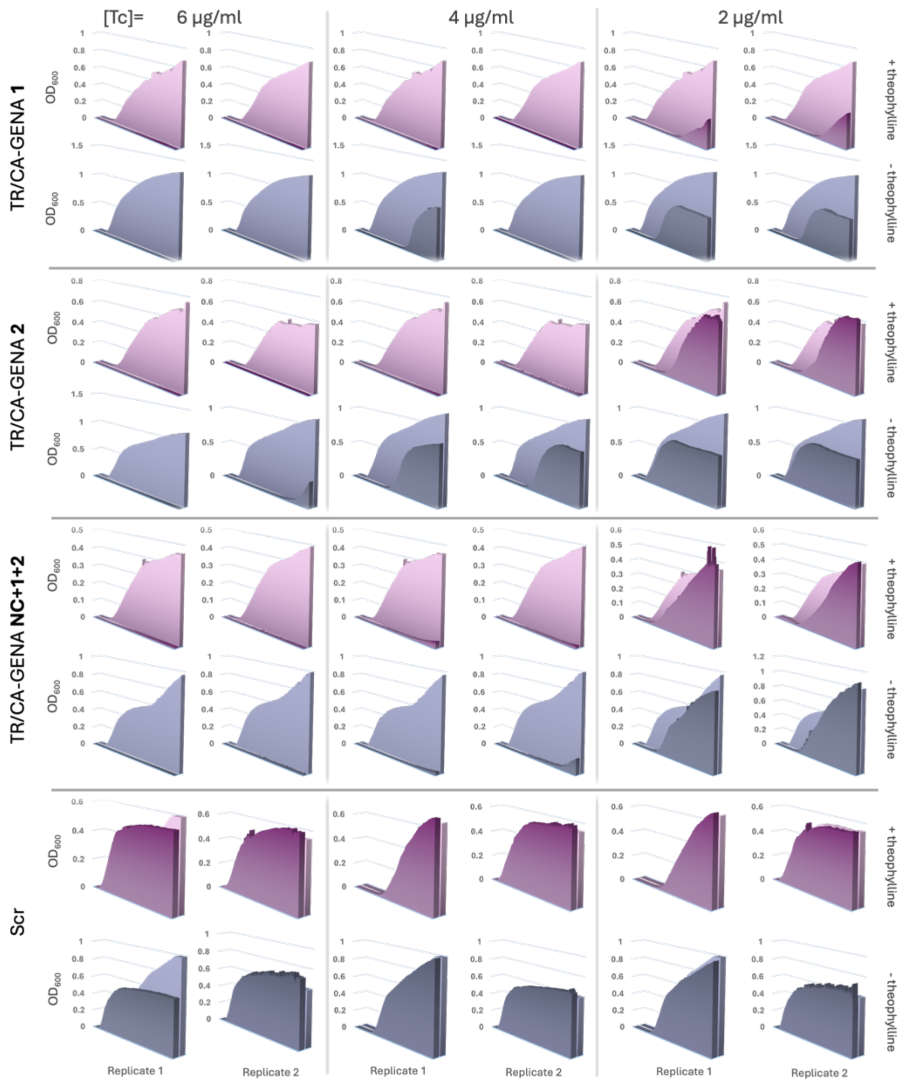

**Figure S1. Growth Curve Analysis of Tetracycline Resistance for Different Clones (part 2):**

**Supplementary Note 2 – Additional Commentary on Table S1 and Figure S1**

While the main text focuses on the phenotypic data obtained in *E. coli* DH5α strains, the results with *E. coli* TOP10 were analyzed in a more concise manner. To provide further clarity, we include here additional notes and interpretations to complement the data presented in Table S1 and Figure S1 regarding experiments conducted in the Top10 background.

In our experiments with *E. coli* TOP10 strains, we consistently observed across two independent experimental batches that clones carrying TR/CA-GENA NC exhibited distinct growth rate patterns depending on the presence or absence of theophylline (Table S1 and Figure S1). Specifically, in the presence of theophylline, growth inhibition occurred at lower tetracycline (Tc) concentrations (compared to assays in the absence of theophylline), indicating a theophylline-dependent reduction in Tc resistance. A similar trend was observed for clones carrying both TR/CA-GENA NC and TR/CA-GENA NC+1. For assays conducted at Tc concentrations above 2 μg/mL (2.5 μg/mL, 3 μg/mL, and 3.5 μg/mL), growth was consistently more inhibited in theophylline-positive conditions compared to theophylline-negative. However, at 3.5 μg/mL, Tc toxicity became strong enough to suppress growth equally in both conditions. These results demonstrate that theophylline-driven inhibition of *tetA* expression leads to a reduction in Tc resistance, even in the TOP10 background, further supporting the robustness and reproducibility of the aptazyme system across strains.

Additionally, we conducted nickel (Ni) sensitivity assays with the TOP10 strains. In theophylline-positive conditions, clones carrying TR/CA-GENA NC and TR/CA-GENA NC+1 maintained stable growth up to 1 μg/ml Ni, whereas in - theophylline conditions, growth began to decline at this concentration. This confirms the expected role of *tetA* in Ni resistance, as its aptazyme-mediated inhibition by theophylline impacts positively the strain's ability to tolerate nickel exposure.

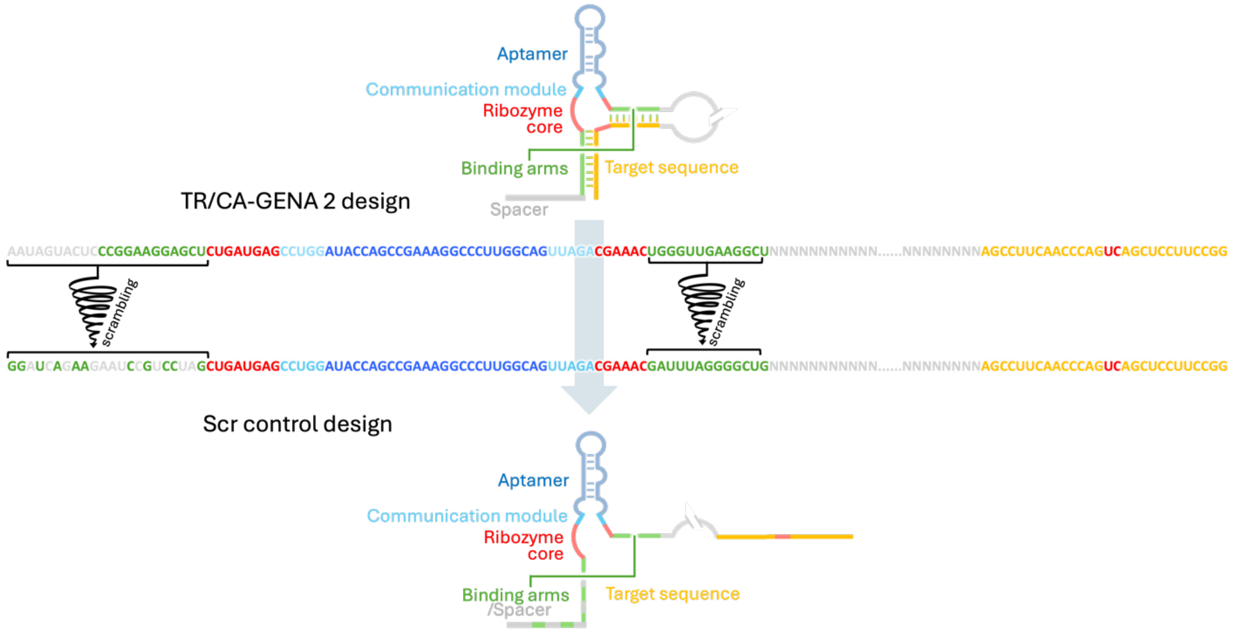

**Figure S2. Scramble Control Design:** The top panel represents the original TR/CA-GENA 3 design, showing how its binding arms (green) can hybridize with the complementary target sequence (yellow). This interaction facilitates ribozyme-mediated cleavage, enabling gene regulation in a ligand-dependent manner. The lower panel illustrates the scramble control design, which was derived by scrambling the hybridizing arms while maintaining the aptamer, communication module, and ribozyme core intact. As a result of this sequence rearrangement, the binding arms can no longer hybridize with the target sequence, disrupting the regulatory function of the aptazyme. This design serves as a negative control, ensuring that observed regulatory effects are specifically due to aptazyme-target hybridization rather than non-specific structural effects.

| | Proximity to target (nt) | Predicted binding strength (kcal/mol) | cleavage (%) | in vivo performance (%) | in vivo leakage activity (%) |
|---|---|---|---|---|---|
| TR/CA-GENA **NC** | **870** | **68** | **0** | **12** | **8** |
| TR/CA-GENA **1** | **753** | **61** | **11** | **87** | **58** |
| TR/CA-GENA **2** | **353** | **75** | **40** | **30** | **31** |

Highest value for a given category

Lowest value for a given category

**Table S2: Comparative summary of in vitro and in vivo parameters for TR/CA-GENA aptazyme designs.** Color-coded summary of TR/CA-GENA aptazyme designs based on structural and functional parameters. Binding strength was estimated in silico using ViennaRNA Fold [1] by simulating aptazyme/target interactions separated by a 19 U linker, with minimum free energy (ΔG) used as a proxy. In vivo repression and leakage values were selected from conditions with [Tc] = 2 µg/mL, either with theophylline (for repression) or without (for leakage), as these revealed the greatest differences between designs. Stronger colors indicate higher values within each category.

### Supplementary Note 3 – Additional Commentary on Table S2:

Monitorable parameters partially predict in vivo aptazyme performance**:** To investigate how structural and functional properties of our aptazyme designs relate to their regulatory performance, we assembled a comparative table (Table 1) that compiles key measurable parameters for each construct. These include values obtained from in vitro cleavage assays, in silico structural modeling, and in vivo gene regulation assays. The purpose of this table is to assess whether these parameters correlate with each other in a predictable way—under the assumption that a design with a high value in one category (e.g., strong binding or high cleavage rate) would likely also perform well in others. If this were the case, each row of the table would be characterized by a consistent color intensity across columns, reflecting a shared trend. However, this is not uniformly observed.

Some encouraging patterns do emerge—for instance, the control design TR/CA-GENA NC, which shows no in vitro cleavage due to a disrupted catalytic core, also performs poorly in vivo. Meanwhile, both cleaving designs (TR/CA-GENA 1 and 2) show significantly better repression activity, confirming the functional importance of cleavage. Binding strength also appears to influence cleavage efficiency in vitro, with stronger binding predicting more cleavage, as seen with TR/CA-GENA 2. Yet, this correlation breaks down when comparing in vivo outcomes: TR/CA-GENA 2, despite its higher in vitro activity, underperforms compared to TR/CA-GENA 1 in living cells. This divergence suggests the presence of additional in vivo factors—such as RNA accessibility or ribosomal interference. For instance, ribosome stalling at slow-translating codons or strong ribosome traffic along coding regions can reduce the accessibility of regulatory RNAs by sterically masking their target sites or unfolding structured RNA elements like aptazymes or ribozymes [2,3]. Similar effects have been described for antisense and small RNAs, which must access mRNA regions before ribosomes occupy them to be effective [4]. These translation-dependent constraints may partially explain why high in vitro cleavage does not always translate into strong in vivo repression.

Additionally, the relatively strong in vivo repression seen with TR/CA-GENA 1 is accompanied by higher leakage in the absence of ligand, consistent with its elevated regulatory capacity. Lastly, proximity to the start codon does not appear to strongly influence regulatory performance under our conditions.

Complementary In Silico and Experimental Approaches Supporting Comparative Table S2:

To generate a comparative overview of aptazyme design features and performance, we compiled five parameters into a color-coded matrix (Table S2). In vivo repression efficiency was quantified at Tc 2 µg/mL in the presence of theophylline, a condition that yielded the greatest dynamic range between constructs. In vivo leakage activity was assessed at the same tetracycline concentration but in the absence of theophylline, representing unintended regulatory effects. In vitro cleavage rates were derived from quantification of radiolabeled fragments following 1-hour incubation with each aptazyme variant.

Binding strength was estimated in silico using the ViennaRNA Fold software. To simulate the aptazyme’s binding capacity independently from tertiary interference, we modeled each construct hybridized to its target RNA with a 19-thymidine spacer separating the aptazyme sequence from the target region. This flexible linker was introduced to avoid artificial structural constraints during energy minimization. The calculated minimum free energy (ΔG) of the resulting RNA–RNA duplex was used as a proxy for binding strength. Target site proximity was measured in nucleotides from the aptazyme binding site to the start codon of the reporter gene. All numerical values were color-coded according to relative magnitude within each column to highlight trends and facilitate visual interpretation.

| Name | Sequence | Notes/function |
|---|---|---|
| EY303N | GAAAGGGAGACCCTGATGAGCCTGGATACCAGCCGAAAGGCCCTTGGCAGTTAGACGAAACGCACTGG | Template sequence for all insert generation. |
| EY450’ | CAATAGAATTCCTGCAGGGTCCTCGCCGAAAATCTGATGAGCCTGGATACCAGCC | Forward primer used to generate DNA insert for TR/CA-GENA 1 design, includes EcoRI and PstI restriction sites. ; compatible with EY303N template. |
| EY451’ N | CTTATGAATTCCCGGGTCCCTGATTTGGCAGCGCTCTGGGTTTCGTCTAACTGCCAAGGGCC | Reverse primer paired with EY450’. Includes EcoRI restriction site. |
| EY452 | GCCCGGGAATAGTACTCCCGGAAGGAGCTCTGATGAGCCTGGATACCAGCC | Forward primer for generating DNA insert for TR/CA-GENA 2 design. Includes XmaI restriction site; compatible with EY303N template. |
| EY453 | GCCCGGGAGCCTTCAACCCAGTTTCGTCTAACTGCCAAGGGCC | Reverse primer paired with EY452. Includes XmaI restriction site. |
| EY454 | GTCTGCAGCGGCGGCCAAAGCGGTCGCTGATGAGCCTGGATACCAGCC | Forward primer for generating DNA insert for TR/CA-GENA 3 design. Includes PstI restriction sites; compatible with EY303N template. |
| EY455 | GTCTGCAGCTGTGCTGGTATTGCGCCGTTCTCGGAGCACTGTTTCGTCTAACTGCCAAGGGC | Reverse primer paired with EY454. Includes PstI restriction sites |
| EY805 | GATCTGAATTCGGATCAGAAGAATCCGTCCTAGCTGATGAGCCTGGATACCAGCC | Forward primer used to generate DNA insert for Scr-control design, includes EcoRI restriction sites. ; compatible with EY303N template. |
| EY806 | GATCTGAATTCCAGCCCCTAAATCGTTTCGTCTAACTGCCAAGGGCC | Reverse primer paired with EY805. Includes EcoRI restriction site. |
| EY1000 | CTAATACGACTCACTATAGGATAATGCCAGCGTAGGGA | Forward primer containing T7 promoter for generating DNA templates. Templates generated are transcribed in vitro into RNA containing cleavage sites for all aptazyme designs (paired with EY1001 and EY2001). (using pLacthiMONtetA as template) |
| EY1001 | GGCGTGCAAGATTCCGAATA | Reverse primer paired with EY1000 for amplifying DNA templates containing cleavage sites for all aptazyme designs. When paired specifically with EY2000, it amplifies a shorter DNA fragment containing only the cleavage sites of TR/CA-GENA 1 and 2 |
| EY2000 | CTAATACGACTCACTATACCTTGCATGCACCATTCCTT | Forward primer paired with EY1001 containing T7 promoter for generating DNA templates. Templates generated are transcribed in vitro into RNA containing the cleavage sites of TR/CA-GENA1 and 2. (using pLacthiMONtetA as template) |
| EY2001 | GAAGCGAGCAGGACTGGG | Reverse primer paired with EY1000 for amplifying DNA templates containing only the cleavage sites of TR/CA-GENA 3 |
| EY900 | TAATACGACTCACTATAGCTTCCGGCTCGTATGTTGTG | Forward primer containing the T7 promoter. When paired with EY801, it amplifies from each plasmid construct used as a template a DNA fragment containing specifically the aptazyme sequence present in that plasmid. (e.g., using the plasmid encoding TR/CA-GENA 1 will yield an amplified fragment containing the TR/CA-GENA 1 aptazyme.) |
| EY801 | ATTTCTCAGCCTTCACGCAG | Reverse primer paired with EY900 |

**Table S3. List of oligonucleotides and their usage in the project**

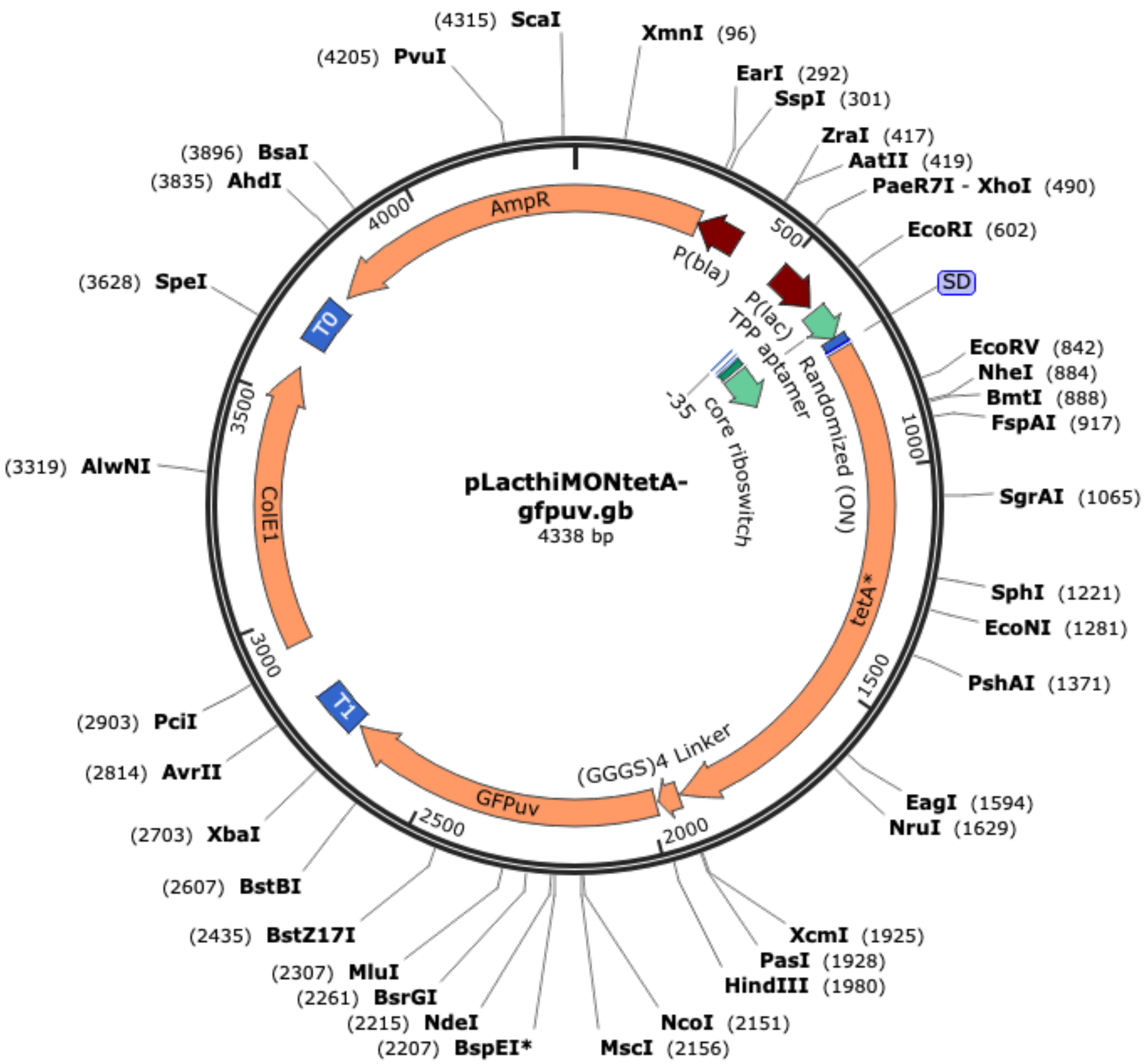

**Figure S3. pLacthiMONtetA-gfpuv map**
